# RACK1 associates with RNA-binding proteins Vigilin and SERBP1 to control dengue virus replication

**DOI:** 10.1101/2021.10.28.466260

**Authors:** Alexis Brugier, Mohamed-Lamine Hafirrassou, Marie Pourcelot, Morgane Baldaccini, Laurine Couture, Vasiliya Kril, Beate M. Kümmerer, Sarah Gallois-Montbrun, Lucie Bonnet-Madin, Sébastien Pfeffer, Pierre -Olivier Vidalain, Constance Delaugerre, Laurent Meertens, Ali Amara

**Author notes:** Corresponding author **Materials & Correspondence** Correspondence and material requests should be addressed to Dr Ali Amara. Institut de Génétique Humaine, Laboratoire de Virologie Moléculaire, CNRS-Université de Montpellier, 34000 Montpellier, France.

## Abstract

Dengue virus (DENV), a re-emerging virus transmitted by *Aedes* mosquitoes, causes severe pathogenesis in humans. No effective treatment is available against this virus. We recently identified the scaffold protein RACK1 as a component of the DENV replication complex, a macromolecular complex essential for viral genome amplification. Here, we show that RACK1 is important for DENV infection. RACK1 mediates DENV replication through binding to the 40S ribosomal subunit. Mass spectrometry analysis of RACK1 partners coupled to a loss-of-function screen identified the RNA binding proteins Vigilin and SERBP1 as DENV host dependency factors. Vigilin and SERBP1 interact with DENV viral RNA (vRNA), forming a ternary complex with RACK1 to mediate viral replication. Overall, our results indicate that RACK1 recruits Vigilin and SERBP1, linking the DENV vRNA to the translation machinery for optimal translation and replication.

## Introduction

Dengue virus (DENV) belongs to the genus *Flavivirus* of the family *Flaviviridae*, which includes important emerging and reemerging viruses such as West Nile virus (WNV), yellow fever virus (YFV), Zika virus (ZIKV), and tick-borne encephalitis virus (TBEV) ^1^. DENV is transmitted to human by an *Aedes* mosquitoe bite and may lead to a variety of diseases ranging from mild fever to lethal dengue hemorrhagic fever and dengue shock syndrome ^2^. Recent estimations indicate that half of the world’s population lives in areas where dengue fever is endemic ^3^, with 100 million symptomatic infections including 500,000 cases of severe manifestations of the disease per year ^4^. There are currently no approved antiviral therapies against DENV, although a promising inhibitor targeting the viral NS3-NS4B interaction was recently described ^5^. Conversely, the recently approved tetravalent lived-attenuated vaccine showed disappointing efficacy ^6,7^.

DENV is an enveloped virus containing a positive-stranded RNA genome of ∼11-kb. Upon entry into the host cell, the viral genome is released in the cytoplasm and translated by the host machinery into a large polyprotein precursor that is processed by host and viral proteases. Co-and post-translational processing gives rise to three structural proteins, [C (core), prM (precursor of the M protein) and E (envelope) glycoproteins] which form the viral particle and seven non-structural proteins (NS) called NS1, NS2A, NS2B, NS3, NS4A, NS4B and NS5^8^ that play central roles in viral genome replication, assembly and modulation of innate immune responses^9^. Like other flaviviruses, DENV genome replication takes place within virus-induced vesicles (Ve) derived from invaginations of the endoplasmic reticulum (ER) membrane ^10,11^. These structures consist of 90 nm-wide vesicles containing a ± 11 nm pore allowing exchanges between the Ve lumen and the cytosol ^11^. Within the Ve, viral NS proteins, viral RNA (vRNA), and some host factors assemble to form the viral replication complex (RC) that is essential for viral RNA synthesis. We have recently purified the DENV RC in human cells, using a tagged DENV subgenomic replicon, and determined its composition by mass spectrometry ^12^. Our study provided an unprecedented mapping of the DENV RC host interactome and identified cellular modules exploited by DENV during active replication. By combining these proteomics data with gene silencing experiments, we identified a set of Host Dependency Factors (HDFs) that critically impact DENV infection and established an important role for Receptor for Activated C-Kinase 1 (RACK1) in DENV vRNA amplification ^12^, which was recently confirmed by others ^13^.

RACK1 is a core component of the 40S ribosomal subunit ^14,15^, containing seven WD40 domains that mediate protein / protein interactions ^16,17^. RACK1 is a scaffold protein ^18,19^ described to interact with many cellular pathways such as Sarcoma (Src) tyrosine kinase ^20,21^, cAMP/PKA ^22^ or receptor tyrosine kinase ^23^. Ribosomal RACK1 has also been shown to be involved in the association of mRNAs with polysomes ^24^, the recruitment and phosphorylation of translational initiation factors ^25–27^ and in quality control during translation ^28^. The non-ribosomal form of RACK1 is involved in innate immunity, by recruiting the PP2A phosphatase ^29^ or by targeting the VISA/TRAF complexes ^30^ and participates in the assembly and activation of the NLRP3 inflammasome ^31^. To date, only one proteomic study aiming to identify RACK1 cofactors has been performed in Drosphila S2 cells ^32^. RACK1 cellular partners in human cells are largely unknown.

Several viruses depend on RACK1 to complete their infectious cycle ^31–35^. For instance, RACK1 is involved in IRES-mediated translation of viruses possessing a type I IRES such as cricket paralysis virus or hepatitis C virus ^33^. RACK1 also contribute to poxvirus infection through a ribosome customization mechanism. Indeed, poxviruses trigger the phosphorylation of the serine 278 of RACK1 ^34^ to promote the selective translation of viral RNAs.

In this work, we have investigated the function of RACK1 during DENV life cycle. We performed the first interactome of RACK1 in human cells. Functional studies revealed that RACK1 forms with the RNA binding proteins Vigilin and SERBP1 a ternary complex that binds viral RNA to regulate DENV genome amplification.

## Results and discussion

### RACK1 interaction with the 40S ribosomal subunit is required for DENV infection

To confirm the role of RACK1 in DENV infection, we challenged parental and RACK1 knockout (RACK1^KO^) HAP1 cells with DENV2-16681 particles at different multiplicity of infections (m.o.i) and measured viral infection by quantifying the percentage of cells expressing the DENV antigen PrM. In agreement with our previous studies ^12^, DENV infection was severely impaired in HAP1 cells lacking RACK1 (Fig. 1 a, b). Importantly, *trans*-complementation of the HAP1 RACK1 ^KO^ cells with a plasmid encoding human RACK1 rescued cell susceptibility to DENV infection (Fig. 1 a, b), ruling out off-target effects and demonstrating that RACK1 is an important host factor for DENV.

**Figure 1:**
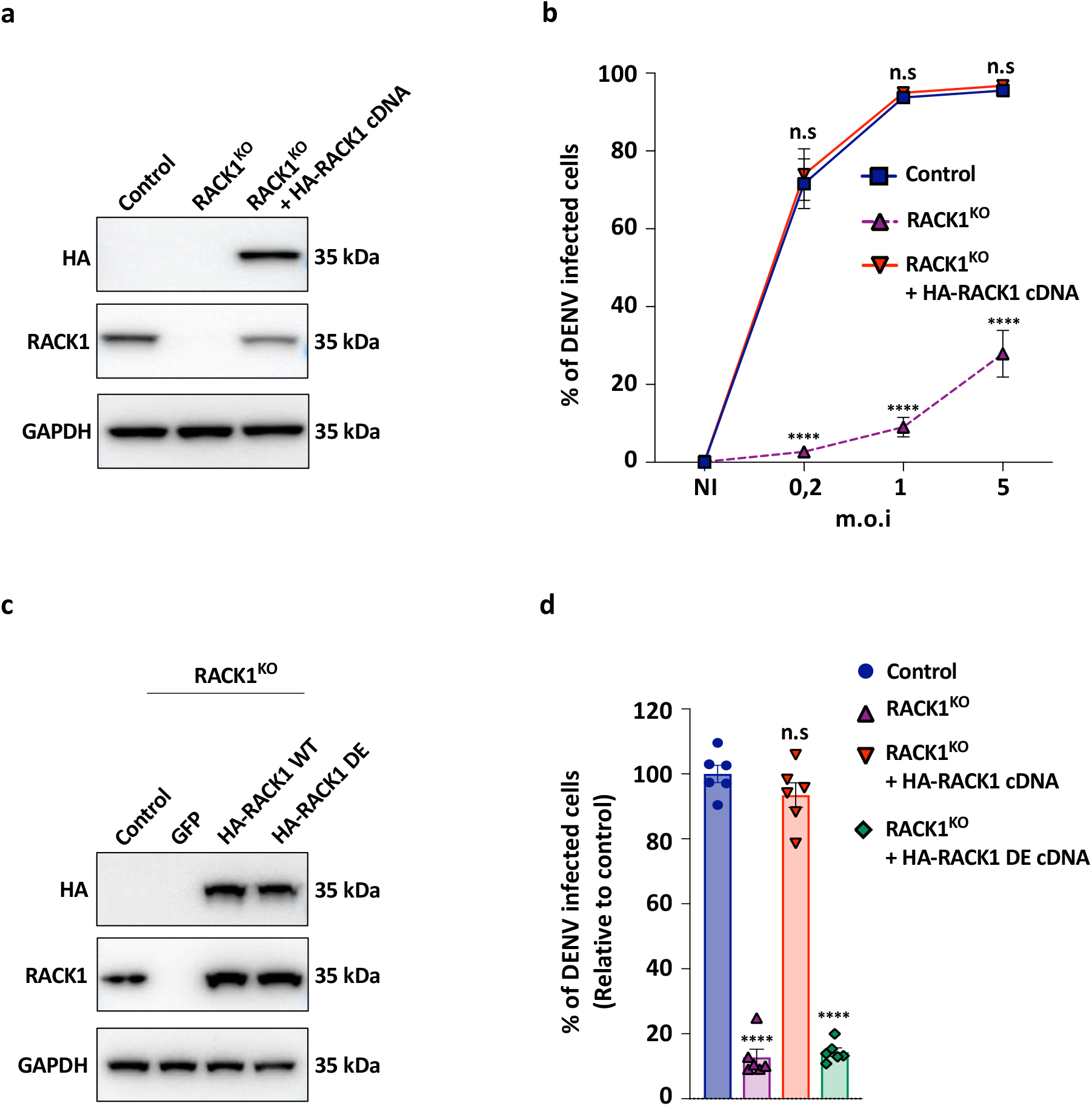
The interaction between RACK1 and the 40S ribosome is required for DENV infection. **(a)** Western Blot analysis of RACK1 expression in control, RACK1^KO^ and RACK1^KO^ HAP1 cells trans-complemented with a HA-RACK1 cDNA. Cell lysates were probed with the indicated antibodies. Representative western blot of n=3 technically independent experiments. **(b)** Role of RACK1 in DENV infection. Controls, RACK1^KO^ or RACK1^KO^ cells transcomplemented with a cDNA encoding WT HA-RACK1 were infected at different m.o.i with DENV2-16681. Levels of infection were determined by flow cytometry using the 2H2 prM mAb at 48 hpi. Data shown are mean +/-s.e.m of 4 independent experiments performed in duplicate. Significance was calculated using two-way ANOVA with Dunnett’s multiple comparison test **(c)** Western Blot analysis of RACK1 expression in RACK1^KO^ HAP1 transcomplemented with cDNA encoding WT HA-RACK1 or the HA-RACK1 D/E mutant. Cells lysates were probed with the indicated antibodies. Representative western blot of 3 independent experiments. **(d)** Impact of RACK1 association to the 40S subunit of the ribosome in DENV infection. Control, RACK1^KO^ and RACK1^KO^ HAP1 cells trans-complemented with cDNA encoding WT HA-RACK1 or the HA-RACK1 DE mutant were infected at m.o.i 1 with DENV2-16681 and harvested at 48 hpi. Levels of infection were determined by flow cytometry as described above. Data shown are mean +/-s.e.m of 3 independent experiments performed in duplicate. Significance was calculated using one-way ANOVA with Dunnett’s multiple comparison test. ns, not significant; ****, *P* < 0.0001

RACK1 is a component of the 40S subunit of the ribosome and is located in near the mRNA exit channel ^17^. To test whether DENV infection requires RACK1 association with the 40S ribosome, we *trans*-complemented RACK1 ^KO^ cells with a RACK1 mutant defective for ribosome-binding (RACK1R36D/K38E, DE mutant) ^34,35^. The RACK1 DE mutant which displayed WT expression level and was unable to associated with polysomes (supplementary Fig.1), as previously described ^36^ and, importantly failed to rescue DENV2-16681 infection (Fig. 1 c, d). These results indicate that the interaction with the 40S ribosomal subunit is important for RACK1 proviral function.

### Mapping the RACK1 interactome

Because RACK1 is a scaffold protein, we hypothesized that its proviral activity may rely on its ability to recruit host proteins near the ribosome for optimal translation. To characterize the RACK1 interactome in mammalian cells, we transfected 293T cells with a plasmid encoding an HA-tagged version of human RACK1. We pulled-down RACK-1 and its binding partners using HA beads and eluated purified proteins with HA peptide according to the experimental procedure that we recently described^12^. Immunoprecipitated proteins were separated by SDS-PAGE, visualized by silver-staining, and subjected to mass spectrometry (MS) analysis (Fig. 2a, supplementary Fig. 2 a, b). By analyzing the raw AP-MS dataset with SAINT express and MiST softwares ^37^, we identified 135 high confidence host factors that co-purified with RACK1 and showed a SAINT express score >0.8 (Table 1). Next, we analyzed the list of 135 high-confidence interactors with DAVID 6.8 to identify statistical enrichments for specific Gene Ontology (GO) terms from the “cellular component” (CC) annotation ^38,39^ (Fig. 2b) and built the corresponding interaction network using Cytoscape 3.4.0 ^40^ (Fig.2c). The 135 RACK1-interacting proteins were clustered into functional modules using enriched GO terms as a guideline and literature mining (Fig. 2c). As expected, the RACK1 interactome was significantly enriched in proteins associated to ribosome/polysome and mRNA translation (Rps3, EIF3, eIF4G, eIF4J), stress granules (G3BP2, LARP1), P-Bodies (Ago1 and 2) and RNA splicing factors (HNRNPA2B1, U2AF2) (Fig. 2c).

**Figure 2:**
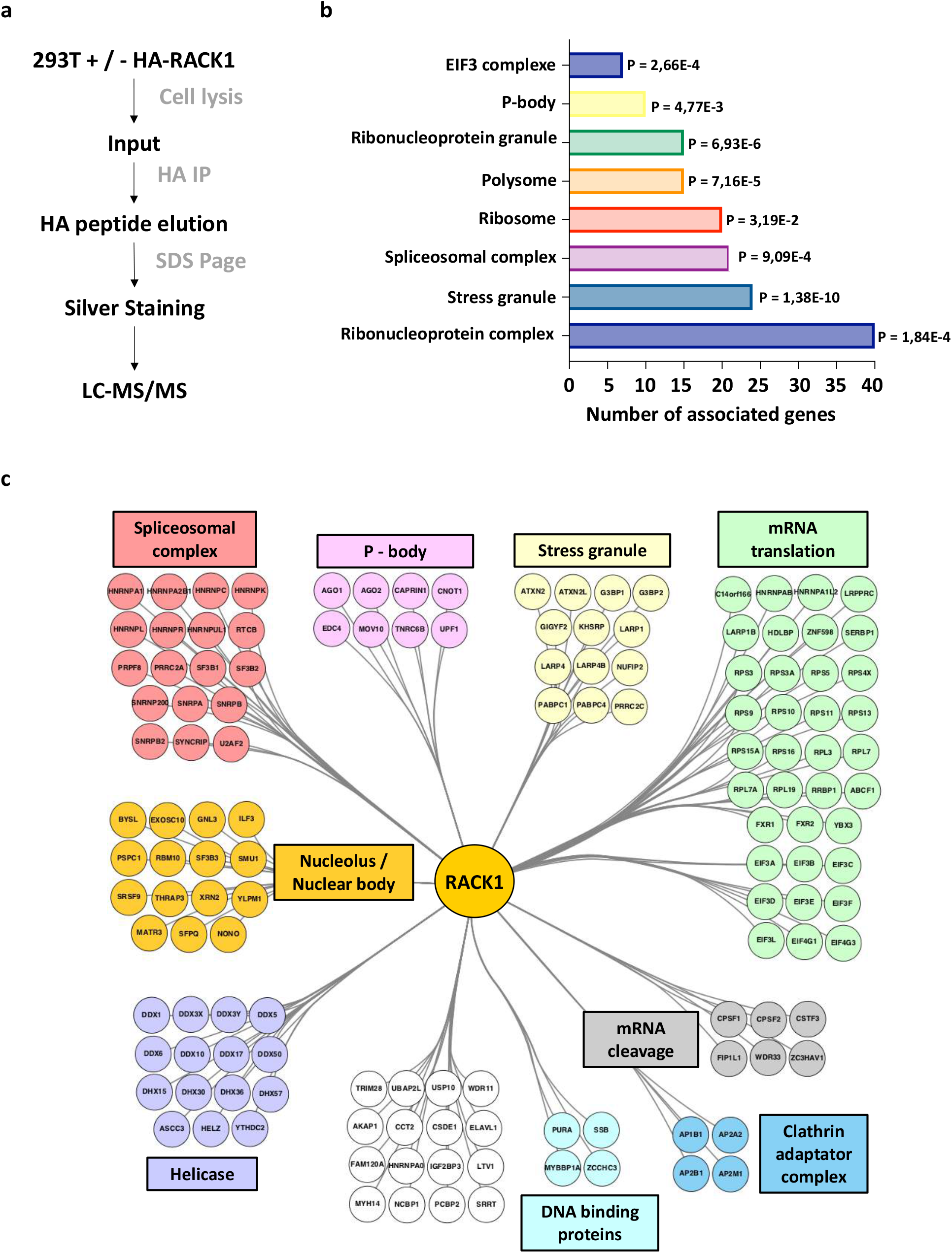
Global map of the RACK1 interactome in human cells. **(a)** Experimental scheme of our RACK1 immunoprecipitation approach. 293T cells expressing RACK1 or HA-RACK1 were lysed, and extracts were purified with anti-HA-coated beads before SDS-PAGE and mass spectrometry (MS) analysis. **(b)** Histogram indicating statistical enrichment for specific biological processes (BP) and cellular components (CC), determined by Gene Ontology (GO) analysis. **(c)** Interaction network of RACK1-associated proteins identified by MS in 293T cells. Proteins were clustered into functional modules using enriched GO terms as a guideline and manual mining of literature. Representative network of n=3 independent experiments showing similar results.

### Vigilin, SERBP1 and ZNF598 are DENV host dependency factors

To pinpoint the function of the RACK-1 binding partners during DENV infection, we silenced by RNA interference (RNAi) the expression of the 49 highest ranked hit with an average peptide count >28, (Fig. 3a, supplementary Fig. 2c) and determined the consequences on viral infection (Fig. 3a, supplementary Fig. 3a, Table 2). Four proteins, namely HNRNPA2B1, Vigilin, SERBP1 and ZNF598 whose silencing decreased infection by at least 50% without affecting cell viability in the two cell lines were considered for further investigations (Fig. 3a, supplementary Fig. 3a, Table 2) These factors are RNA-Binding Proteins (RBP) involved in RNA splicing (HNRNPA2B1) ^41^ or translation regulation (Vigilin, SERBP1, ZNF598) ^28,42,43^. HNRNPA2B1 was already described to interact with the 3’UTR part of the virus ^44^. Because HNRNPA2B1 is a nuclear protein ^45^, it was not further considered in our study. Vigilin is a multiple K-homology (KH) domain protein implicated in translation regulation and lipidic metabolism ^24,43,46^. This protein was recently described to bind the DENV RNA and, in association with the ribosomal-binding protein 1 (RRBP1), to facilitate viral RNA translation and replication ^47^. However, how this protein interacts with RACK1 to regulate DENV infection is still unknown. SERBP1 is a RACK1 cofactor ^48^ that is located at the entry channel of ribosomes ^49^ and enhances translation by promoting the association of mRNAs with polysomes ^42^. SERBP1 was also described to interact with DENV RNA, however its role in DENV replication remains unclear ^50^. Finally, ZNF598 is an E3 ubiquitin-protein ligase known to interact with RACK1 and playing a key role in the ribosome quality control ^28^. ZNF598 was also described to play a role in innate immunity ^51^, however its role in DENV infection is unknow.

**Figure 3:**
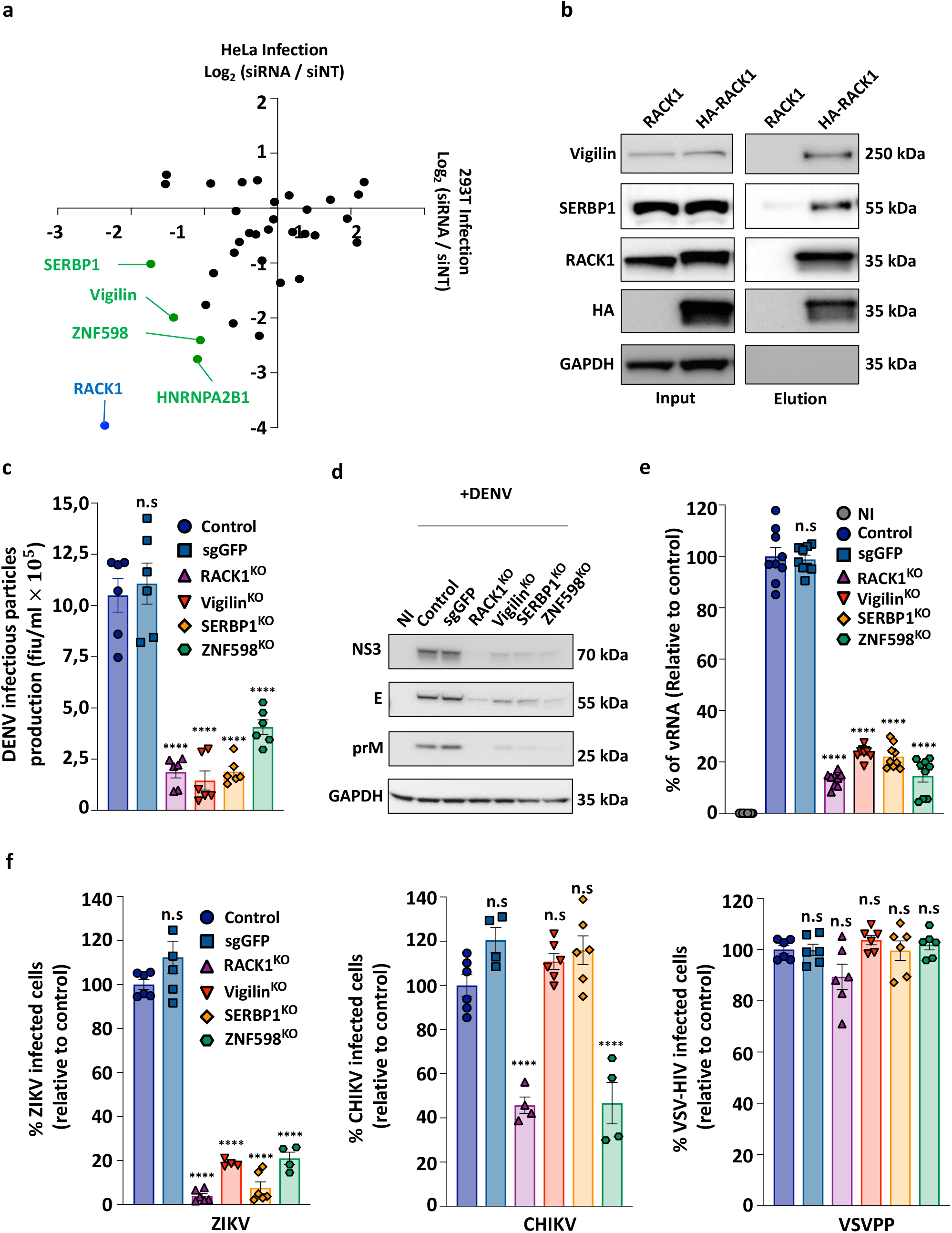
Vigilin, SERBP1 and ZNF598 are DENV host dependency factors. **(a)** Host dependency factors (HDFs) found in our RNAi screen. Data shown are representative of 3 independent experiments. Host dependency factors are marked in green. Positive control (siRNA pool targeting RACK1) is highlighted in blue. **(b)** Validation of the interaction between RACK1 and endogenous Vigilin or SERBP1 in 293T cells by immunoprecipitation. Cell extracts from 293T cells expressing RACK1 or HA-RACK1 were subjected to affinity-purification using anti-HA beads and interacting proteins were revealed by western blot. Data shown are representative of 3 independent experiments. **(c-e)** Impact of RACK1/Vigilin/SERBP1/ZNF598 gene editing on DENV infectious cycle in HAP1 cells. The indicated cells were infected for 48 hrs at m.o.i 1 with DENV2-16681. **(c)** Supernatants from infected cells were harvested, then titered by flow cytometry on Vero cells and expressed as FIU/ml. FIU, FACS Infectious Unit. **(d)** Infection was assessed by immunoblot using anti-NS3, anti-prM and anti-E DENV mAb. Data shown are representative of 3 independent experiments. **(e)** Levels of infection were assessed by quantification of DENV vRNA by qRT-PCR using NS3 primers. **(c and d)** Data shown are mean +/-s.e.m of 3 independent experiments performed in duplicate. Significance was calculated using a two-tailed Student’s t test. **(f)** The indicated cells were infected with ZIKV HD78 at m.o.i 2 (left), CHIKV 21 at m.o.i 2 (middle), VSV-pp at m.o.i 2 (right). Levels of infection were determined by flow cytometry at 48 hpi. Data shown are mean +/-s.e.m of at least 2 independent experiments performed in duplicate. Significance was calculated using one-way ANOVA with Dunnett’s multiple comparison test.. ns: not significant; ****: *P* < 0.0001

We first confirmed that endogenous Vigilin, ZNF598 and SERBP1 proteins co-immunoprecipitated with HA-RACK1 ectopically expressed in 293T cells (Fig. 3b). Next we validated the requirement of Vigilin, SERBP1 and ZNF598 using two approaches. On one hand, we found that knocking-down by RNA interference Vigilin, SERBP1or ZNF598 significantly impaired DENV infection of primary human fibroblasts which are DENV target cells (Supplementary Fig. 3b, c). On the other hand, we use the CRISPRCas9 technology to edit the corresponding genes in HAP1 cells (Vigilin ^KO^, SERBP1 ^KO^, ZNF598 ^KO^). Gene editing and knock-out generation were confirmed by genomic DNA sequencing (Supplementary Fig. 3d) and western blot analysis (Supplementary Fig. 3e), respectively. In agreement with our previous findings, lack of RACK1, Vigilin, SERBP1 and ZNF598 expression had no impact on cell growth and viability as assessed by quantification of ATP levels in culture wells at different timepoints (Supplementary Fig. 3f). HAP1 cells lacking Vigilin, SERBP1 or ZNF598 expression were poorly permissive to DENV infection as shown by the quantification of viral progeny in supernatants of infected cells (Fig. 3c), western blot analysis of the DENV protein expression (NS3, E, PrM) (Fig. 3d), and quantification of the viral RNA (Fig. 3e). Parental (Control) and HAP1 cells transfected with a non-specific gRNA (sgGFP) were used as negative controls (Fig. 3) while RACK1 ^KO^ HAP1 cells as a positive control (Fig. 3). We then investigated if these phenotypes were specific to DENV2-16681 or could be observed with other flaviviruses. We found that Vigilin, SERBP1 and ZNF598 mediate infection by other DENV serotypes (Supplemental Fig. 3g), as well as by Zika virus (ZIKV), a related flavivirus (Fig. 3f). In contrast, infections by the alphavirus Chikungunya virus (CHIKV) or vesicular stomatitis virus G protein (VSV-G)-pseudotyped Human Immunodeficiency Virus (VSVpp), were unaffected in Viligin ^KO^ and SERBP1 ^KO^ cells (Fig. 3f). CHIKV infection but not VSVpp was significantly reduced in RACK1 ^KO^ and ZNF598 ^KO^ cells (Fig. 3f). Altogether, our data indicate that Vigilin, SERBP1 and ZNF598 are important host factors for DENV. ZNF598 is required for DENV and CHIKV infection while Vigilin and SERBP1 are exclusively exploited by DENV and other related flaviviruses.

### Vigilin and SERBP1 regulate DENV translation and replication

To determine whether Vigilin and SERBP1 impact initial vRNA translation or amplification, Vigilin ^KO^ and SERPB1 ^KO^ cells were challenged with DENV2 *Renilla* luciferase (Luc) reporter virus (DV-R2A) through a time-course experiment to monitor the kinetic of viral infection (Fig. 4a). RACK1 ^KO^ cells were used as a positive control.

**Figure 4:**
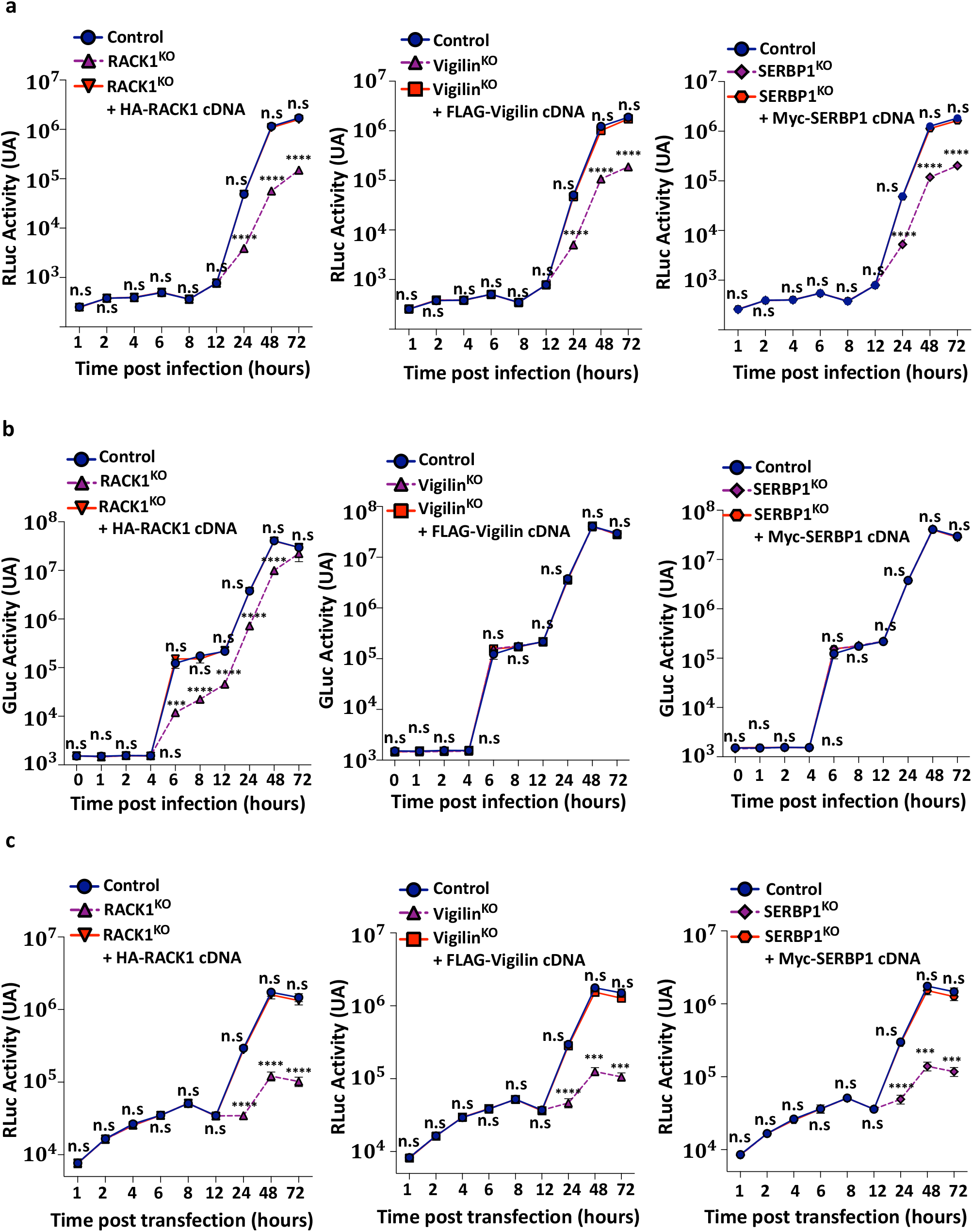
Vigilin and SERBP1 regulate DENV translation and replication. **(a)** The indicated cells were infected at m.o.i 1 with DENV-Luc. At indicated time points Renilla luciferase activity reflecting RNA translation (1 to 8 hpi) and replication (12 to 72 hpi) was measured. Data shown are mean +/-s.e.m of 3 independent experiments performed in triplicate. two-way ANOVA with Dunnett’s multiple comparison test. **(b)** Indicated cells were infected at m.o.i 1 with CHIKV-Luc. Gaussia luciferase activity was monitored at the indicated time points. Data shown are mean +/-s.e.m of 3 independent experiments performed in triplicate. Significance was calculated using two-way ANOVA with Dunnett’s multiple comparison test. **(c)** Impact of RACK1/Vigilin/SERBP1 KO on DENV life cycle in HAP1 cells transfected with a DENV replicon RNA expressing Renilla luciferase. Renilla luciferase activity was monitored at the indicated time point. Data shown are mean +/-s.e.m of 3 independent experiments performed in triplicate. Significance was calculated using two-way ANOVA with Dunnett’s multiple comparison test. ns: not significant; ****: *P* < 0.0001.

A small peak of the Luc activity was detected at 6 hr post-infection, reflecting the initial translation of the incoming vRNA. This was followed by a marked increase in Luc activity due to vRNA amplification (translation and replication) (Fig. 4a). Depletion of RACK1, Vigilin and SERBP1 had no impact on initial translation step but strongly impaired DENV vRNA amplification at later time-points (Fig. 4a). Importantly, viral genome replication was completely restored in KO cells transduced with RACK1, SERBP1 or Vigilin cDNAs (Fig. 4a and supplementary Fig. 4a). Consistent with Fig. 3f, CHIKV expressing the Gaussia luciferase replicated as efficiently in Vigilin or SERBP1 ^KO^ cells than control cells while its replication in RACK1 ^KO^ was impaired (Fig. 4b). To assess further the effect of Vigilin and SERBP1 on DENV vRNA replication, we used a *Renilla* luciferase (Rluc) reporter sub-genomic replicon (sgDVR2A). This latter is a self-replicating DENV RNA containing a large in-frame deletion in the structural genes and represents a useful tool to exclusively monitor DENV translation and RNA amplification. Control, Vigilin ^KO^, SERBP1 ^KO^, and RACK1 ^KO^ HAP1 cells were transfected with the *in vitro*-transcribed DENVR2A sub-genomic RNA and vRNA replication was monitored over time by quantifying the Rluc activity in infected cell lysates (Fig. 4c). Depletion of RACK1, Vigilin or SERBP1 had no impact during the early phase of DENV RNA translation. At 12h post-transfection, the RLuc signal increased over time in control cells, while a strong reduction was observed (more than 10-fold reduction at 48 hpi) in Vigilin ^KO^ and SERBP1 ^KO^ cells (Fig. 4c). The RLuc signal was restored in Vigilin ^KO^ or SERBP1 ^KO^ trans-complemented with their corresponding cDNAs (Fig. 4c).

Vigilin has been previously shown to mediate, in association with the host factor RRBP1, the stability of DENV vRNA ^47^. Since SERBP1 also binds the DENV RNA ^50^, we reasoned that it might play a similar role. To assess this hypothesis RACK1^KO^, Vigilin^KO^ or SERBP1^KO^ HAP1 cells where challenged with DENV followed by treatment with MK0608 to inhibit viral replication ^45^. Then, we monitored the decay of the vRNA overtime by northern blot analysis using a probe that targets the DENV 3’UTR (supplementary data Fig. 4b). We observed that the levels of the DENV genomic RNA was similar in control, RACK1 ^KO^ and SERBP1 ^KO^ HAP1 cells up to 24 h after MK0608 treatment (supplementary data Fig. 4b). Surprisingly, lack of Vigilin expression had a very mild effect on DENV RNA stability (supplementary data Fig. 4b). Together, these results show that RACK1, Vigilin and SERBP1 promote viral replication without a major impact on the stability of DENV vRNA.

### Vigilin and SERBP1 interactions with RACK1 are important for DENV infection

Scp160p and Asc1p, the yeast homologs of Vigilin and RACK1 respectively, have been shown to interact each other ^24^. This interaction is thought to promote translation of specific mRNAs linked to Scp160p by mediating their association with polysomes ^24^. Because Vigilin is very well-conserved amongst different species, a similar interaction with RACK1 might occur in mammalian cells. Having established that Vigilin and SERBP1 do not have a major influence on the stability of the vRNA, we hypothesized that their proviral effect might be linked to their interaction with RACK1. Previous studies showed that Scp160p interacts with Asc1p via the KH 13 and 14 domains located in its C-terminal region ^52^ while SERBP1 interacts directly with RACK1 through a motif (aa 354 to 474) which contains the RGG domain ^48^ (supplementary data Fig. 5a). On the bases of these observations, we generated the corresponding deletion mutants of Flag-tagged Vigilin (Flag Vigilin Mut) and Myc-tagged SERBP1 (Myc SERPB1 Mut) (supplementary data Fig. 5a)and test their ability to interact with RACK1 (Fig. 5a, supplementary data Fig. 5). Pull-down experiments showed that RACK1 binds both WT FLAG Vigilin or WT Myc SERBP1 ectopically expressed in HEK-293T cells (Fig. 5a). In contrast, RACK1 failed to associate with mutant forms of Vigilin and SERBP1 (Fig. 5a). Using an RNA-IP assay, we showed that Vigilin Mut and SERBP1 Mut bound the DENV vRNA as the same extent as their WT counterparts (Fig. 5b and supplementary Fig. 5b). Finally, infection studies showed that expression of Mut Vigilin or Mut SERBP1 in Vigilin ^KO^ or SERBP1 ^KO^ cells, respectively, did not restore DENV2-16681 infection in contrast to their WT counterparts (Fig. 5c, supplementary Fig. 5c). Together, these data indicate that the ability of Vigilin and SERBP1 to bind RACK1 but not the vRNA is required for DENV infection.

**Figure 5:**
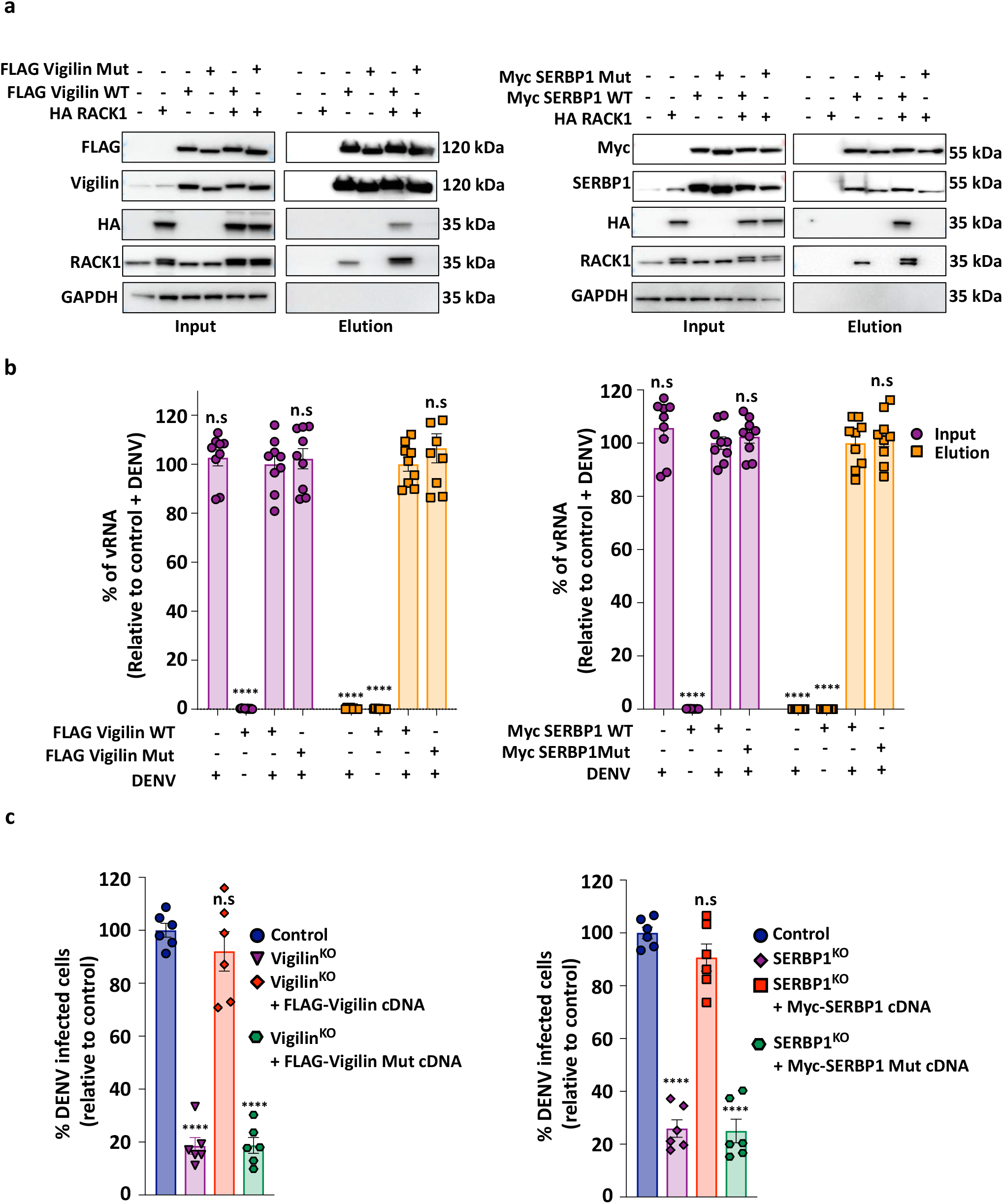
Vigilin and SERBP1 interaction with RACK1 is important for DENV infection. **(a)** Evaluation of FLAG-Vigilin mutant (left) or Myc-SERBP1 mutant (right) interaction with RACK1. Cell extracts from 293T expressing Wt or mutated form of Vigilin and SERBP1 were subjected to affinity-purification using anti-FLAG or -Myc coated beads, respectively. Input and eluates were resolved by SDS-PAGE and interacting proteins were revealed by western blot using corresponding antibodies. Representative western blot of 3 independent experiments. **(b)** RIP analysis of the interaction of Vigilin (WT and Mut) and SERBP1 (WT and Mut) with the DENV vRNA. Cells were infected at m.o.i 1 by DENV2-16681 and harvested 48 hpi. Tagged-proteins were immunoprecipitated after UV crosslink at 254 nm using anti-FLAG or -Myc coated beads. The amount of vRNA in the input (in purple) and the elution fractions (in orange) were determined by RT-qPCR using DENV2 specific primers. Data shown are mean +/-s.e.m of 3 independent experiments performed in triplicate. Significance was calculated using a one-way ANOVA with Dunnett’s multiple comparison test **(c)** DENV infection in HAP1 cells expressing Vigilin (WT or Mut) and SERBP1 (WT or Mut). The indicated cells were infected at m.o.i 1 with DENV2-16681. Levels of infection were determined by flow cytometry at 48 hpi. Data shown are mean +/-s.e.m of 3 technically independent experiments performed in duplicate.. Significance was calculated using one-way ANOVA with Dunnett’s multiple comparison test. ns: not significant; ****: *P* < 0.0001.

## Conclusions

In this study, we performed the first RACK1 interactome in human cells and identified Vigilin and SERBP1 as host factors for DENV infection. Both are RNA-binding proteins that interact with the DENV RNA and regulate viral replication. Importantly, our data suggest that Vigilin, SERBP1 and RACK1 form a ternary complex important for DENV RNA amplification. The proviral function of RACK1 dependents on its association with the 40S ribosomal subunit. Furthermore, mutants of SERBP1 or Vigilin that lost their ability to interact with RACK1 were unable to support infection. Overall, our results provide new insights into the molecular mechanisms of DENV replication and indicate that RACK1 acts as a binding platform at the surface of the 40S ribosomal subunit to recruit Vigilin and SERBP1, which may therefore function as linkers between the viral RNA and the translation machinery to selectively amplify DENV genome. Strategies that interfere with RACK1-ribosome association or disturb the RACK1-Vigilin -SERBP1 complex may represent new ways to combat DENV-induced disease.

## Methods

### Cell lines

HAP1 cells (Horizon Discovery), and HAP1 RACK1 ^KO^ (provided by Dr Gabriele Fuchs; University at Albany) were cultured in IMDM supplemented with 10% fetal bovine serum (FBS), 1% penicillin–streptomycin, 1% GlutaMAX and 25 mM HEPES. HEK293T (ATCC), Vero E6 (ATCC), BHK-21 (ATCC), HeLa (ATCC) cells were cultured in DMEM supplemented with 10% FBS, 1% penicillin–streptomycin, 1% GlutaMAX and 25 mM HEPES. Fibroblast BJ-5ta cells (ATCC) were cultured according to the manufacturer’s instructions. A final concentration of 50 μM of MK0608 was used in this study. All cell lines were cultured at 37°C and 5% CO2.

### Virus strains and replicons

DENV1-KDH0026A (gift from L. Lambrechts, Pasteur Institute, Paris), DENV2-16681 (Thaïland/16681/84), DENV4 (H241) and ZIKV HD78788 were propagated in mosquito AP61 cell monolayers with limited cell passages. DENV2 Rluc reporter virus (DVR2A) wasprovided by Ralf Bartenschlager (University of Heidelberg). The CHIKV Luc reporter virus was described previously ^53^. To generate infectious virus, capped viral RNAs were generated from the NotI-linearized plasmids using a mMessage mMACHINE T7 Transcription Kit (Thermo Fisher Scientific) according to the manufacturer’s instructions. RNAs were purified (see RNA IP protocol), resuspended in Dnase/Rnase free water, aliquoted and stored at **−**80 °C until used. 30 μg of purified RNAs were transfected in BHK21 cells using lipofectamine 3000 reagent. Supernatants were collected 72 hrs later and used for viral propagation on Vero E6 cells. For all of the viral stocks used in flow cytometry experiments, viruses were purified through a 20% sucrose cushion by ultracentrifugation at 80,000g for 2 h at °C. Pellets were resuspended in HNE1X pH 7.4 (HEPES 5 mM, NaCl 150 mM, EDTA 0.1 mM), aliquoted and stored at −80 °C. Viral stock titers were determined on Vero E6 cells by plaque-forming assay and were expressed as plaque-forming units (PFU) per ml. Virus stocks were also determined by flow cytometry as described ^54^. Vero E6 cells were incubated 1 hour with 100 μl of tenfold serial dilutions of viral stocks. The inoculum was then replaced with 500 μl of culture medium and the percentage of infected cells was quantified by flow cytometry using the 2H2 anti-PrM mAb at 8 h after infection. Viral titers were calculated and expressed as FIU per ml: Titer = (average percentage of infection) x (number of cells in well) x (dilution factor) / (ml of inoculum added to cells).

To establish a DENV replicon plasmid, based on the infectious DENV2-16681 cDNA clone, the region encoding the structural proteins was mostly deleted and replaced by a cassette encoding ubiquitin - Renilla luciferase - foot-and-mouth disease virus (FMDV) 2A. DENV replicon RNA was generated as previously described ^11^. Infection or replication was determined by measuring the luciferase activity using TriStar LB942 microplate reader (Berthold Technologies). RFP-expressing lentiviral vector pseudotyped with vesicular stomatitis virus glycoprotein G (VSV-G) were generated by transfecting HEK293FT cells with pNL4.3 Luc RFP ΔEnv, psPAX2 and pVSV-G (4:3:1 ratio) using lipofectamine 3000. Supernatants were harvested 48 h after transfection, cleared by centrifugation, filtered, and frozen at -80°C.

### Antibodies and reagents

All antibodies and reagents are listed in Table 3

### Polysome profiling

2 × 10^8^ of indicated cells were incubated with 100 μg/mL of cycloheximide (CHX) for 10 min at 37°C and washed twice with cold PBS + 100 μg/mL CHX. Cells were pelleted by centrifugation at 4 °C, 300 × g for 10 min and washed once with cold PBS + 100 μg/mL CHX. The pellet was resuspended in 2 ml Lysis Buffer (10 mM Tris-HCl pH7.5; 100 mM KCl; 10 mM Magnesium acetate; 1% Triton X100; 2 mM DTT) containing 100 μg/ml CHX. Cells were pulverized by adding glass beads and vortexed for 5 min at 4°C. Cells debris were removed by centrifugation at 4 °C, 3,000 × rpm for 10 min and the supernatant was transferred to a 2 ml cryovial. The determination of polysome concentration was done by spectrophotometric estimation, based on the fact that ribosomes are ribonucleoprotein particles. Supernatant was quickly flash-frozen in liquid nitrogen and stored in a −80 °C freezer. The supernatant was loaded on a 10– 50 % sucrose gradient (31% sucrose; 50 mM Tris-acetate pH 7.6; 50 mM NH_4_Cl; 12 mM MgCl 2; 1 mM DTT) and spinned for 3 h at 39,000 rpm, 4 °C, in an SW41 swing-out rotor. The gradient was fractionated by hand and analyzed by immunoblotting.

### Mass spectrometry analysis

HAP1 cells (5 × 10^8^), expressing either the WT or the HA-tagged RACK1 proteins, were lysed in Pierce IP lysis buffer (Thermo Scientific) in the presence of Halt protease inhibitor cocktail (Thermo Scientific) for 30 min at 4°C and then cleared by centrifugation for 30 min at 6,000 × g. Supernatants were incubated overnight at 4°C with anti-HA magnetic beads. Beads were washed three times with B015 buffer (20 mM Tris-HCl pH 7.4, 150 mM NaCl, 5 mM MgCl2, 10% glycerol, 0.5 mM EDTA, 0.05% Triton, 0.1% Tween-20) and immune complexes were eluted twice with HA peptide (400 mg/mL) 30 min at room temperature (RT). Eluates were concentrated on a Pierce Concentrator, PES 10K, and stored at - 20°C until used. A total of three co-affinity purifications and MS analysis experiments were performed with the HA-tagged RACK1 protein or the untagged RACK1 protein as a control in 293T cells. Samples were analyzed at Taplin Biological Mass Spectrometry Facility (Harvard Medical School). Briefly, concentrated eluates issued from immunopurification of endogenous and RACK1-HA-tagged protein are separated on 10% Tris-glycine SDS-PAGE gels (Invitrogen), and stained with Imperial Protein Stain (Thermo Fisher). Individual regions of the gel were cut into 1 mm^3^ pieces and subjected to a modified in-gel trypsin digestion procedure ^55^. Peptides were desalted and subjected to a nano-scale reverse-phase HPLC ^56^. Eluted peptides were then subjected to electrospray ionization and then MS/MS analysis into an LTQ Orbitrap Velos Pro ion-trap mass spectrometer (Thermo Fisher Scientific, Waltham, MA). Peptides were detected, isolated, and fragmented to produce a tandem mass spectrum of specific fragment ions for each peptide. Peptide sequences were determined by matching protein databases with the acquired fragmentation pattern by the Sequest software program (Thermo Fisher Scientific, Waltham, MA) ^57^. All databases include a reversed version of all the sequences, and the data were filtered to < 2% peptide false discovery rate.

### Network analysis

The AP-MS dataset was analyzed with SAINTexpress and MIST software ^37^. Of the 1671 proteins selected in our pipeline, 193/1671 showed a probability score > 0.80 with SAINTexpress and 135/193 showed an AveragePeptide Count > 10. This list of 135 host proteins was analyzed with DAVID 6.8 to identify statistical enrichments for specific GO terms from the ‘‘cellular component’’ (CC) annotation ^38,39^. The interaction network was built using Cytoscape 3.4.0 ^40^, and proteins were clustered into functional modules using enriched GO terms as a guideline and manual curation of literature.

### siRNA Screen Assay

An arrayed ON-TARGETplus SMARTpool siRNA library targeting 49/135 proteins of our RACK1 network, which had an average peptide count > 28 was purchased from Horizon Discovery. To this end, HeLa or 293T cells were transfected with a 30 nM final concentration of siRNA using the Lipofectamine RNAiMax (Life Technologies). 48 hrs post-transfection, cells were infected with DENV2-16681 at MOI 5. Infection was quantified 48 hrs post infection by flow cytometry and viability by CellTiter-Glo 2.0 Assay (Promega). Two siRNA controls were included in the screen: a non-targeting siRNA used as a reference (siNT) and a siRNA targeting RACK1(siRACK1) as a positive control for host dependency factors HDFs) ^12^. HDFs were defined as factors whose inhibition in both cell types decreases infection by at least 50% compared to siNT and viability by at most 20% of the siNT.

### Gene editing and *trans*-complementation experiments

sgRNA targeting Vigilin, SERBP1 and ZNF598 were designed using the CRISPOR software ^58^. Sequences for all the sgRNAs are listed in the Table 1. The sgRNAs were cloned into the plasmid lentiCRISPR v2 (Addgene) according to the recommendations provided by the members of the Zhang’s laboratory (Broad Institute, Cambridge, MA. HAP1 cells were transiently transfected with the plasmid expressing sgRNAs and selected with puromycin until all mock-transfected cells died. Clonal cell lines were isolated by limiting dilution and assessed by DNA sequencing and immunoblot for gene editing. The human HA-RACK1 WT and HA-RACK1 DE mutant plasmid were provided by the Gabriele Fuchs Lab (University at Albany), the FLAG-tagged Vigilin cDNA was purchased from Genscript (Clone ID: OHu17734) and the Myc-tagged SERBP1 cDNA was purchased from Genscript (Clone ID: OHu26811C). After PCR, amplification products were cloned into a SpeI-NotI (RACK1), NotI-XhoI (Vigilin) or EcoRI-BamHI (SERBP1) digested pLVX-IRES-ZsGreen1 vector. SERBP1 mutant and Vigilin mutant were obtained using the Q5® Site-Directed Mutagenesis Kit (E0554) (NEB) with deletion primers using the WT cDNA in pLVX as template. All primers are listed in Table 1. Lentiviral like particles for transduction were prepared in 293T cells by co-transfecting the plasmid of interest with psPAX2 (from N. Manel’s lab, Curie Institute, Paris) and pCMV-VSV-G at a ratio of 4:3:1 with Lipofectamine 3,000 (Thermo Fisher Scientific). Supernatants were collected 48 h after transfection, centrifugated (750 g, 10 min), filtered using a 0.45μm filter, and purified through a 20% sucrose cushion by ultracentrifugation (80,000 g for 2 h at 4°C). Pellets were resuspended in HNE1X pH 7.4, aliquoted, and stored at **−**80°C. Cells of interest were transduced by spinoculation (750 g for 2 h at 32°C) and sorted for GFP-positive cells by flow cytometry if necessary.

### Flow cytometry analysis

Indicated cells were plated in 24 well plates and infected. At indicated times, cells were trypsinized and fixed with 2% paraformaldehyde (PFA) diluted in PBS for 15 min at room temperature. Cells were incubated for 1 hour at 4°C with 1 μg/ml of 3E4 anti-E2 monoclonal antibody (CHIKV), 2H2 anti-prM monoclonal antibody (mAb) (DENV) or the anti-E protein mAb 4G2 (ZIKV). Antibodies were diluted in permeabilization flow cytometry buffer (PBS supplemented with 5% FBS, 0.5% saponin, 0.1% sodium azide). After washing, cells were incubated with 1 μg/ml of Alexa Fluor 488 or 647-conjugated goat anti-mouse IgG diluted in permeabilization flow cytometry buffer for 30 min at 4°C. Acquisition was performed on Attune NxT Flow Cytometer (Thermo Fisher Scientific) and data were analyzed by FlowJo software (TreeStar).

### Infectious virus yield assay

To assess the release of infectious particles during infection, indicated cells were inoculated for 3 h with DENV2-16681, washed once with PBS and maintained in the culture medium for 48 h. At the indicated time points, supernatants were collected and kept at -80°C. Vero E6 cells were incubated with three-fold serial dilutions of supernatant for 24 h and prM expression was quantified by flow cytometry as previously described ^54^.

### Immunoblots

Cell pellets were lysed in Pierce IP Lysis Buffer (Thermo Fisher Scientific) containing Halt protease and phosphatase inhibitor cocktails (Thermo Fisher Scientific) for 30 min at 4 °C. Equal amounts of protein, determined by DC Protein Assay (BioRad), were prepared in 4X LDS sample buffer (Pierce) containing 25 mM dithiothreitol (DTT) and heated at 95 °C for 5 min. Samples were separated on Bolt 4–12% Bis-Tris gels in Bolt MOPS SDS Running Buffer (Thermo Scientific) and proteins were transferred onto a PVDF membrane (BioRad) using the Power Blotter system (Thermo Fisher Scientific). Membranes were blocked with PBS containing 0.1% Tween-20 and 5% non-fat dry milk and incubated overnight at 4 °C with primary antibodies (HA 1/5,000, RACK1 1/4,000, GAPDH 1/5,000, Vigilin 1/500, SERBP1 1/2,000, NS3 DENV 1/4,000, 2H2 prM DENV 1/4,000, E DENV 1/5,000, FLAG 1/2,000, Myc 1/1,000, Tubulin 1/500, ZNF598 1/10,000, Anti Mouse HRP 1/5,000, Anti Rabbit HRP 1/10,000. Staining was revealed with corresponding horseradish peroxidase (HRP)-coupled secondary antibodies and developed using Super Signal West Dura Extended Duration Substrate (Thermo Fisher Scientific) following the manufacturer’s instructions. The signals were acquired with Fusion Fx camera (VILBERT Lourmat).

### Co-immunoprecipitation assay

Indicated cells were plated in 10 cm dishes (5 × 10^6^) After 24 h, the cells were transfected with a total of 15 μg DNA expression plasmids (7.5 μg of each plasmid in co-transfection assays) using Lipofectamine 3,000 (Thermo Fisher Scientific). After 24 h of transfection, the cells were washed once with PBS, collected, and centrifugated (400g for 5 min). Cell pellets were lysed in Pierce IP Lysis Buffer (Thermo Fisher Scientific) containing Halt protease and phosphatase inhibitor cocktails (Thermo Fisher Scientific) for 30 min at 4 °C. Equal amounts of protein, determined by DC Protein Assay (BioRad), were incubated overnight at 4 °C, with either anti-FLAG magnetic beads, anti-HA magnetic beads or anti-Myc magnetic beads. Beads were washed three times with BO15 buffer (20 mM Tris-HCl pH 7.4, 150 mM NaCl, 5 mM MgCl_2_, 10% glycerol, 0.5 mM EDTA, 0.05% Triton X-100, 0.1% Tween-20) before incubation. The retained complexes were eluted twice with either 3x FLAG peptide (200 μg/ml Sigma-Aldrich), HA peptide (400 μg/ml Roche) or cMyc peptide (200 μg/ml Sigma-Aldrich) for 30 min at RT. Samples were prepared and immunoblotted as described above. For input, 1% of whole-cell lysates was loaded on the gel.

### RNA preparation and quantitative RT-qPCR

Total RNA extraction from the indicated cells was performed using the RNeasy Plus Mini kit (Qiagen). RNA was quantified using a Nanodrop One (Thermo Fisher Scientific) before cDNA amplification. cDNA was prepared from 100 ng total RNA with Maxima First Strand Synthesis Kit (Thermo Fisher Scientific) including an additional step of RNase H treatment after reverse transcription. RT–qPCR was performed using Power Syber green PCR master Mix (Thermo Fisher Scientific) on a Light Cycler 480 (Roche). Quantification was based on the comparative threshold cycle (Ct) method, using GAPDH as endogenous reference control. All primers are listed in Table 1.

### RNA immunoprecipiation (RNA-IP)

Indicated cells (2 × 10 ^6^)were plated in 10 cm dishes, transfected 48 h with the corresponding plasmids using Lipofectamine 3000 and then infected with DENV2-16681 at m.o.i 2. 48 h post infection, culture media was removed, and cells were washed twice with cold PBS. 10 ml of cold PBS were added on the cell before UV cross-link (2000mJ/cm2). Cells were collected and spun 5 min at 4°C 2,000 rpm. Cell pellets were lysed in 1 ml of Pierce IP Lysis Buffer (Thermo Fisher Scientific) containing Halt protease and phosphatase inhibitor cocktails (Thermo Fisher Scientific) + 250 U of RNasin (Promega) for 30 min at 4 °C. 250 U of turbo DNase was added and the lysate was put 30 min at 37°C and centrifugated at 15,000 rpm for 15 min. The supernatant was then collected. The protein of interest was immunoprecipitated and eluted (see co-immunoprecipitation assay). 100 μl of Input and Elution were incubated with 150 μl of proteinase K buffer (117 μl NT-2, 15 μl SDS 10 %, and 18 μl of Proteinase K) 1 h at 56°C and then 750 μl of Trizol Reagent was added. RNA was extracted by phenol chloroform precipitation: 0.2 ml of chloroform per 1 ml of TRIZOL Reagent was added. Samples were vortexed vigorously for 15 seconds and incubated at room temperature for 2 to 3 minutes and then centrifuged at 12,000 x g for 15 minutes at 4°C. Following centrifugation, the upper aqueous phase was transferred carefully without disturbing the interphase into fresh tube. The RNA from the aqueous phase was precipitated by mixing with 0.5 ml of isopropyl alcohol per 1 ml of TRIZOL reagent used for the initial homogenization. Samples was incubated at RT for 10 minutes and centrifuged at 12,000 x g for 10 minutes at 2 to 4°C. The supernatant was removed completely, and the RNA pellet was washed twice with 1 ml of 75% ethanol per 1 ml of TRIZOL Reagent used for the initial homogenization. The samples were mixed by vortexing and centrifuged at 7,500 x g for 5 minutes at 2 to 8°C. The RNA pellet was air dried for 5-10 minutes and then dissolved in RNase free water.

### Cell viability assay

Cell viability and proliferation were assessed using CellTiter-Glo 2.0 assay (Promega) according to the manufacturer’s protocol. Cells (3 × 10 ^4^) were plated in 48 well plates. At the indicated times, 100 μl of CellTiter-Glo reagent were added to each well. After 10 min of incubation, 200 μl from each well was transferred in an opaque 96-well plate (Cellstar, Greiner Bio-One) and luminescence was measured on a TriStar2 LB 942 (Berthold) with 0.1 s integration time.

### RNA stability measurement by high molecular weight northern blot analysis

Indicated cells (1 × 10 ^6^) were plated on a 60-mm dish and infected with DENV2-16681. 48 h post infection, medium was replaced by MK0608 (50 μM final concentration) containing medium to block viral replication. At the indicated time post treatment, cells were washed twice with cold PBS and harvested in TRIzol (Thermo Fisher Scientific). Total RNA extraction was performed as previously described in RIP protocol. The DENV2-specific probe was obtained after PCR amplification of the 3’UTR of the DENV2-16681 infectious clone (from 10,205 to 10,704). Probes were then labeled with α-^32^P dCTP using the Prime-a-gene kit (Promega). For high molecular weight northern blot analysis to detect DENV2 genomic RNA, 5 μg of total RNA were denatured for 5 minutes at 65°C in RNA sample buffer (32% deionized formamide, 4% formaldehyde, 1X MOPS, ethidium bromide 1 μg/μl). Then, RNA loading buffer (50% glycerol, EDTA 1mM, 0.4% bromophenol blue) was added. RNAs were resolved in a 1% agarose gel containing 1X MOPS and 3.7% formaldehyde in 1X MOPS buffer, before being transferred overnight on a nylon Hybond N+ membrane (Cytiva) in a 20X SSC solution (Euromedex). RNAs were UV crosslinked (120 mJoules) with Stratagene Stratalinker 1800 (LabX). Membrane was blocked and hybridized overnight at 42°C using PerfectHyb™ Plus hybridization buffer (Sigma) with the corresponding labeled probe. The day after, the membrane was washed using 2X SSC, 0.1% SDS solution twice at 42°C and 0.1X SSC, 0.1% SDS twice at 50°C before being exposed on an Image Plate (Fujifilm) during 24 h. The plate was revealed using Typhoon™ FLA 7,000 (GE Healthcare). Densitometry analysis of the bands was performed using Image Quant TL 8.1 software (GE healthcare).

### Graphics and statistical analyses

The number of independent experimental replications is indicated in the legends. Graphical representation and statistical analyses of mean and s.e.m. were performed using Prism 8 software (GraphPad Software) as well as Student’s t-test.

## Acknowledgements

This study has received funding from the Fondation pour la Recherche Medicale (grant FRM - EQU202003010193), the French Government’s Investissement d’Avenir program, Laboratoire d’Excellence “Integrative Biology of Emerging Infectious Diseases” (grant n°ANR-10-LABX-62-IBEID), the ANR-15-CE15-00029 ZIKAHOST. A.B was funded by a scholarship from the French Ministry of Research. The authors thank Karim Majzoub, Alessia Zamborlini for critical readings of the manuscript and helpful discussions. The authors are grateful to Ralf Bartenschalger (Heidelberg University, Germany) and Dr Gabriele Fucks (University at Albany, NY, 12222, USA) for providing us with DENV R2A reporter virus and RACK1 knockout cells and plasmids, respectively. Ali Amara dedicates this work to the memory of Professor Jean-Louis Virelizier (Unité d Immunologie Virale, Institut Pasteur, Paris) and Professor Renaud Mahieux (Ecole Normale Supérieure, Lyon, France), who left us during the SARS-CoV-2 epidemic.

## Author’s contributions

A.B., MLH, and A.A conceived the study. A.B., M.L.H., M.P, L.C., L.B.M., C.D., L.M. and A.A. designed the experiments. A.B. and M.L.H. performed the RACK1 interactome and the RNAi screen. P.O.V. provided help in the data analysis. M.P., L.C., L.B.M., and V.K. generated the viruses used in this study and performed infection studies. B.M.K. generated the DENV replicon and provided expertise in viral RNA production. S.P. and M.B. performed the DENV RNA stability experiments. S.G.M. participated in the RNA-IP experiments. A.B. and A.A. wrote the initial manuscript draft, and the other authors contributed to its editing in its final form.

