## Supplemental information for "RACK1 associates with RNA-binding proteins Vigilin and SERBP1 to control dengue virus replication"

**a**

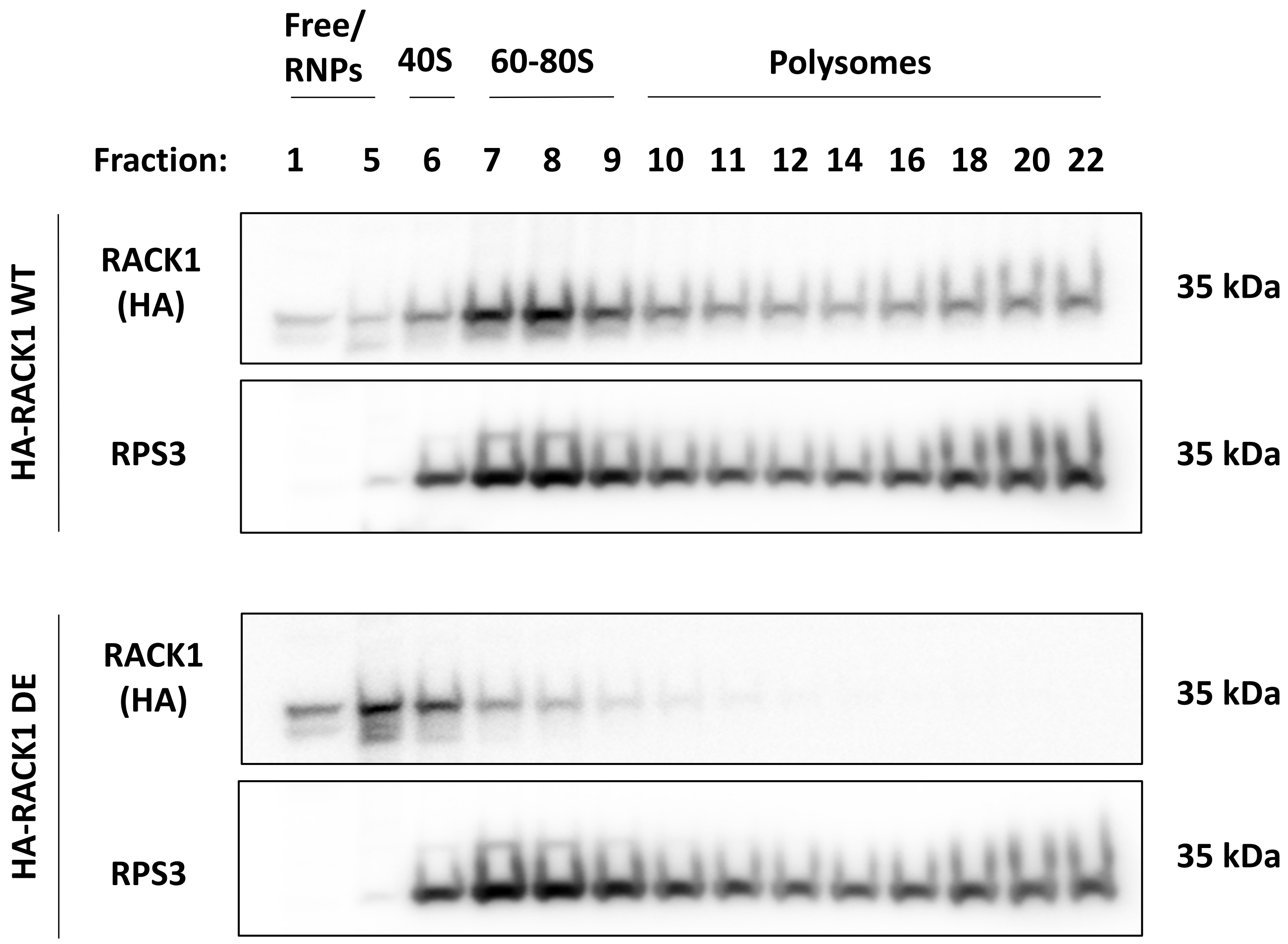


**Supplemental Figure 1**

**Supplemental information related to figure 1**

Characterization of the HA-RACK1 DE mutant by polysome profiling. Fractions from the linear sucrose density gradient from cells overexpressing either HA-RACK1 WT (top) or HA-RACK1 DE mutant (bottom) were immunoblotted with the indicated antibodies. Representative western blot of n=3 independent experiments.

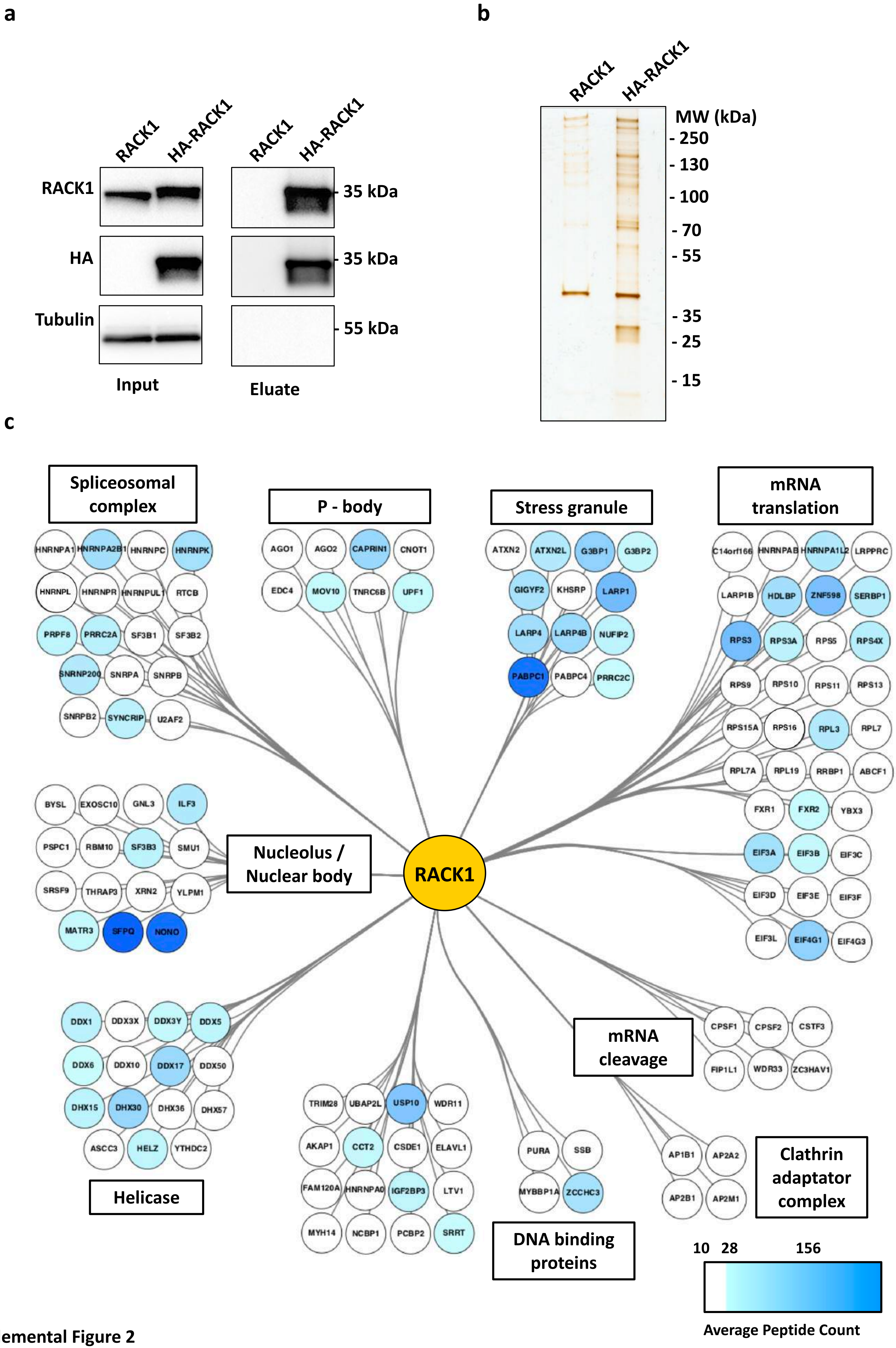


Supplemental Figure 2

**Supplemental information related to figure 2**

**(a-b)**. 293T cells expressing RACK1 or HA-RACK1 were lysed, and extracts were purified with anti-HA-coated beads. **(a)**. Proteins were resolved on SDS-PAGE and analyzed by western blot using indicated antibodies. **(b)**. Proteins were resolved by SDS-PAGE and visualized by silver staining. **(c)** Main hits of the RACK1 proteomic in 293T cells with an average peptide count > 28.

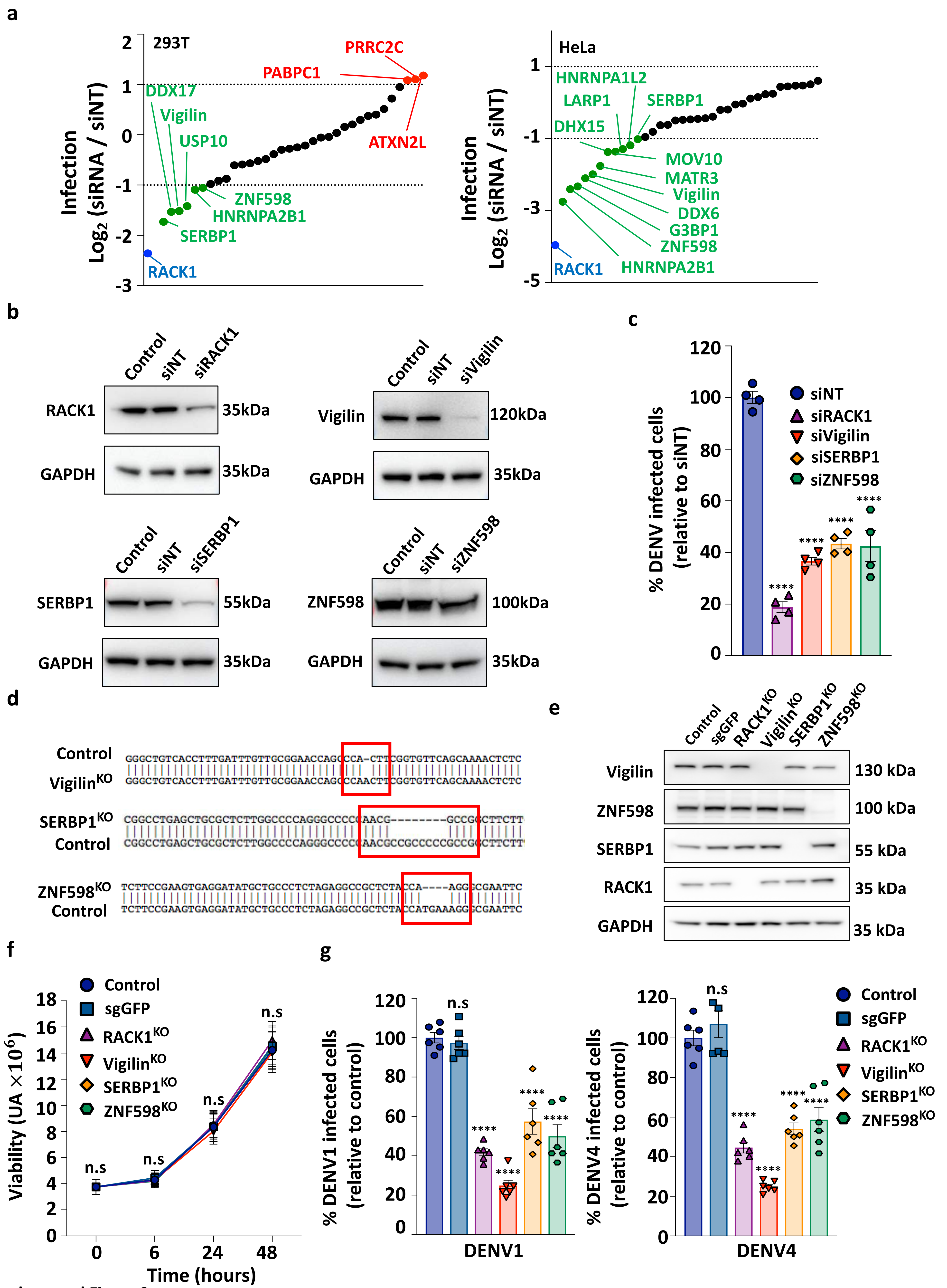


Supplemental Figure 3

**Supplemental information related to figure 3**

**(a)** Host Dependency Factors (HDFs) identified in our RNAi screen in 293T (left) or HeLa (right) cells. Data shown are representative of n=2 independent experiments. Host dependency factors (HDFs) are marked in green and Host Restriction Factors (HRFs) in red. Positive control (siRNA pool targeting RACK1) is highlighted in blue. **(b)** Human primary fibroblasts were transfected with the indicated siRNA pool. RACK1, Vigilin, SERBP1 and ZNF598 expression in siRNA transfected cells was assessed by Western Blot analysis 48 hours post-transfection. **(c)** SiRNA transfected fibroblast described in b were challenged with DENV2-16681 at m.o.i 1 48 hours post-transfection. Levels of infection were determined by flow cytometry using 2H2 mAb at 48 hpi. Data shown are mean +/- s.e.m of 3 independent experiments performed in duplicate. Significance was calculated using one-way ANOVA with Dunnett's multiple comparison test. **(d)** Sanger sequencing of *VIGILIN*, *SERBP1*, *ZNF598* in control and $Vigilin^{KO}$, $SERBP1^{KO}$ or $ZNF598^{KO}$ HAP1 cells respectively. **(e)** Validation of Vigilin, SERBP1, ZNF598 gene editing by Western Blot analysis. Representative western blot of 3 independent experiments. **(f)** Impact of RACK1, Vigilin, SERBP1, ZNF598 gene editing on cell viability at different time point in HAP1 cells by cell titter glow analysis. Data shown are mean +/- s.e.m of 3 independent experiments performed in duplicate. Significance was calculated using two-way ANOVA with Dunnett's multiple comparison test. **(g)** $RACK1^{KO}$, $Vigilin^{KO}$, $SERBP1^{KO}$ or $ZNF598^{KO}$ HAP1 cells infection by different DENV serotypes. Cells were infected at m.o.i 1 with DENV1-KDH or DENV4-H241. Levels of infection were determined by flow cytometry at 48 hpi. Data shown are mean +/- s.e.m of 3 independent experiments performed in duplicate. Significance was calculated using a one-way ANOVA with Dunnett's multiple comparison test. ns: not significant; ****: $P < 0.0001$

**a**

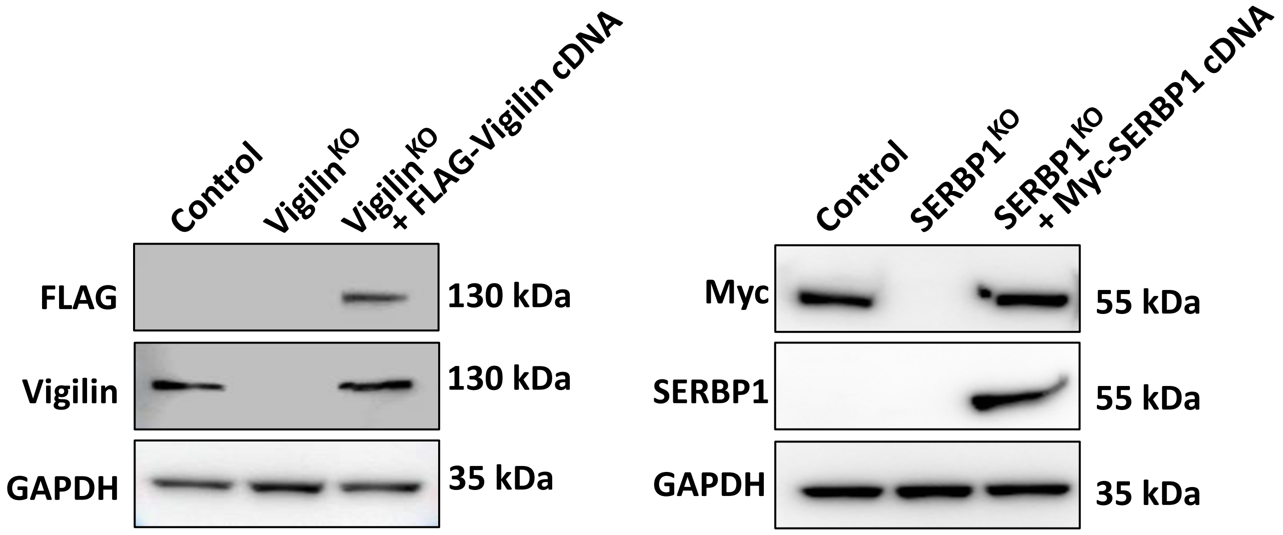


**b**

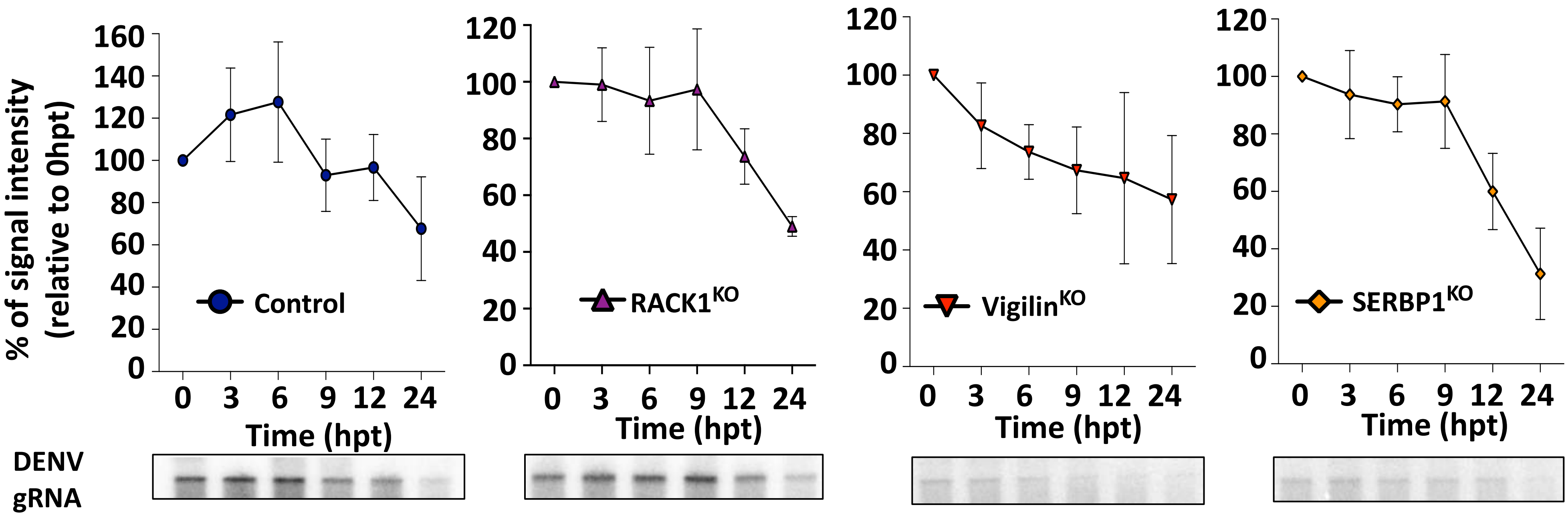


Supplemental Figure 4

**Supplemental information related to figure 4**

**(a)** Immunoblot of Vigilin (left) or SERBP1 (right) expression in Vigilin$^{KO}$ or SERBP1$^{KO}$ HAP1 cells, respectively. Ectopic Vigilin or SERBP1 expression in KO cells stably transduced with FLAG-Vigilin and Myc-SERBP1 is shown as well. Data shown are representative of 3 independent experiments. **(b)** Impact of RACK1, Vigilin, SERBP1 gene editing on DENV genomic RNA stability in HAP1 cells by Northern blot analysis. Cells were infected at m.o.i 1 with DENV2-16681, and vRNA was extracted 48 hpi after MK0608 replication inhibitor treatment during indicated time. Data shown are mean +/- s.e.m of 3 independent experiments performed in triplicate. Data were expressed as percentage relative to the signal monitored at time point 0 hour after MK0608 treatment.

a

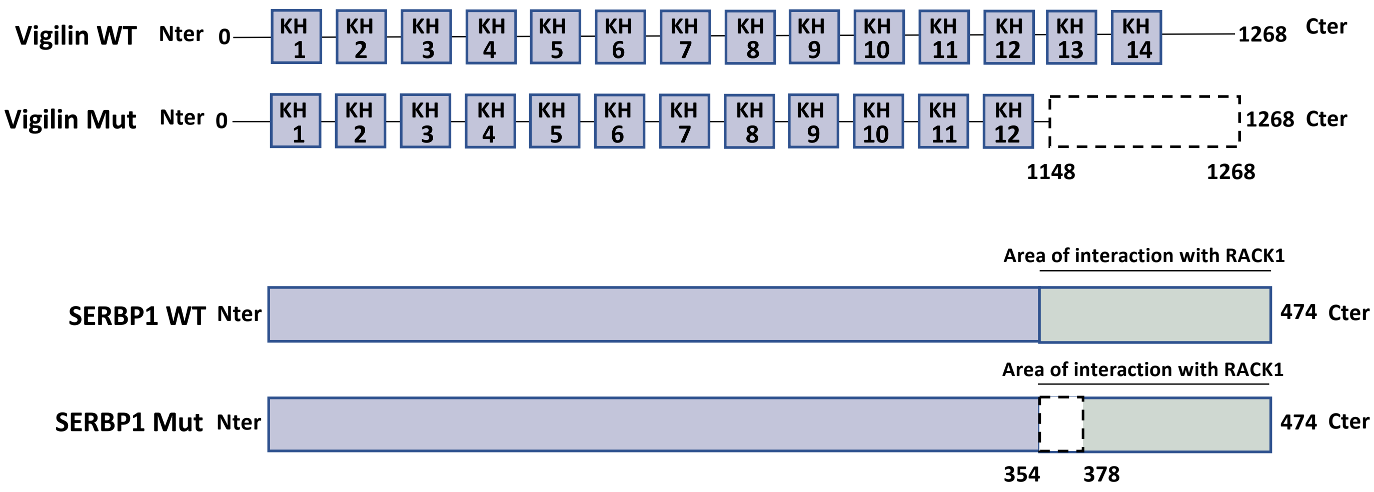


b

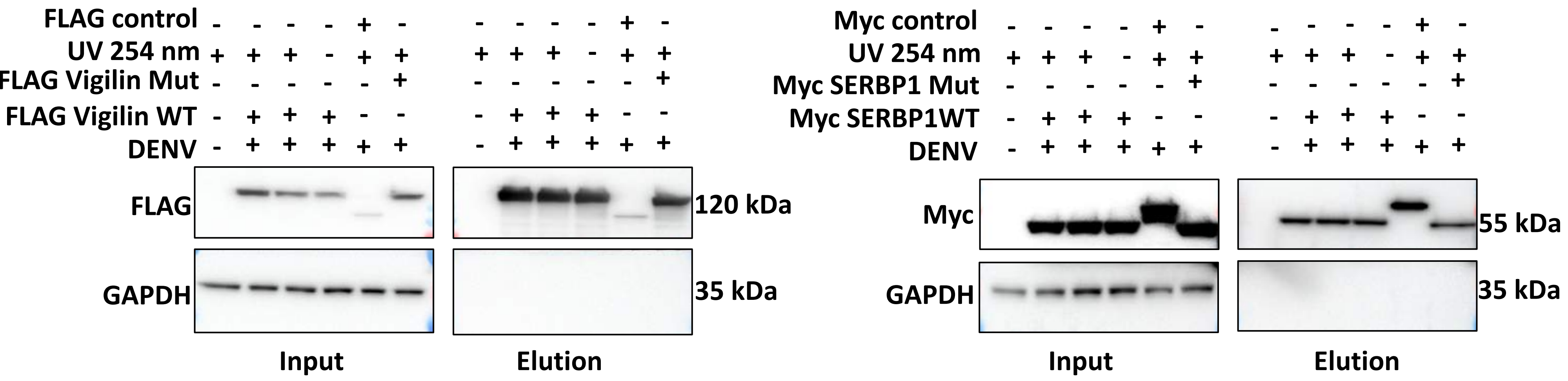


c

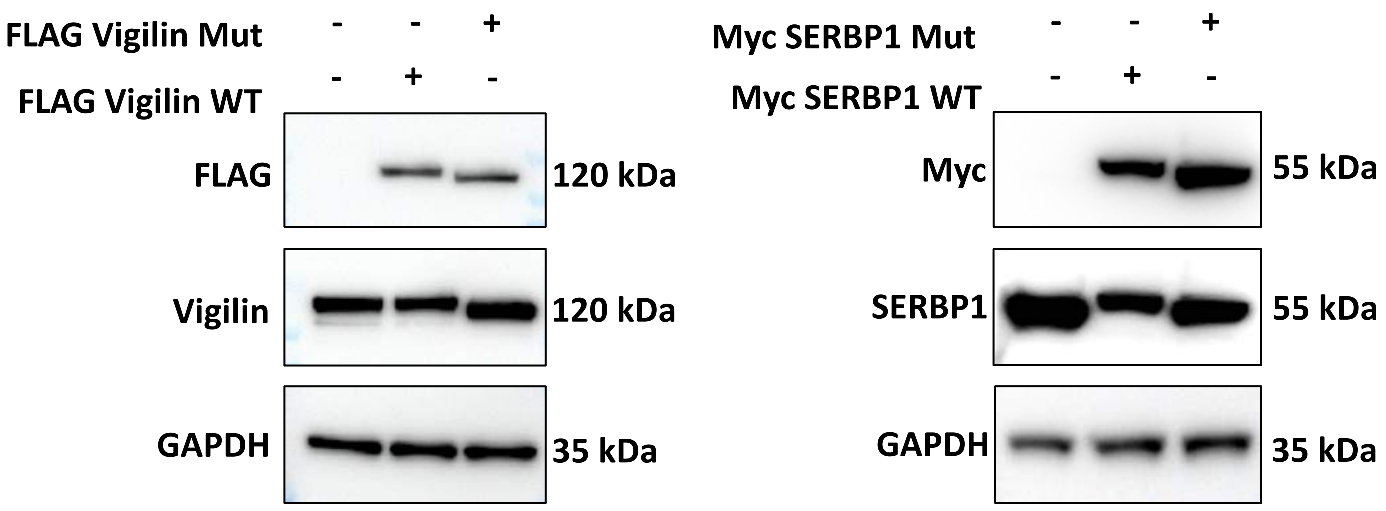


Supplemental Figure 5

**Supplemental information related to Figure 5**

**(a)** Schematic representation of Vigilin mutant (up) and SERBP1 mutant (down) constructs. **(b)** Cell extracts prepared from 293T cells transfected with the indicated plasmids and infected or not by DENV were subjected to affinity-purification using anti-FLAG or -Myc coated beads after UV crosslink at 254 nm. Input and eluates were resolved by SDS-PAGE and interacting proteins were revealed by western blot using corresponding antibodies. Data shown are representative western blot of n=3 independent experiments **(c)** Western blot analysis of FLAG-Vigilin WT and FLAG-Vigilin mutant stably expressed in HAP1 Vigilin $^{KO}$ cells (left panel), or Myc-SERBP1 WT and Myc-SERBP1 expressed in HAP1 SERBP1 $^{KO}$ cells (right panel). Cell lysates were probed with the indicated antibodies. Representative western blot of n=3 independent experiments.
